# Phage Cocktails can Prevent the Evolution of Phage-Resistant *Enterococcus*

**DOI:** 10.1101/2021.01.27.428553

**Authors:** Stephen Wandro, Pooja Ghatbale, Hedieh Attai, Clark Hendrickson, Cyril Samillano, Joy Suh, David T. Pride, Katrine Whiteson

**Affiliations:** Department of Molecular Biology and Biochemistry, University of California, Irvine; Department of Pathology, University of California, San Diego; Department of Medicine, University of California, San Diego

**Author notes:** Address correspondence to Katrine Whiteson, Stephen Wandro and Pooja Ghatbale contributed equally to this work. Author order was determined on the basis of seniority.

## Abstract

Antibiotic resistant *Enterococcus* infections are a major health crisis that requires the development of alternative therapies. Phage therapy could be an alternative to antibiotics and has shown promise in *in vitro* and in early clinical studies. Phage therapy is often deployed as a cocktail of phages, but there is little understanding of how to most effectively combine phages. Here we utilized a collection of 20 *Enterococcus* phages to test principles of phage cocktail design and determine the phenotypic effects of evolving phage resistance in *Enterococcus* isolates that were susceptible or resistant to antibiotics (e.g., Vancomycin Resistant Enterococcus (VRE)). We tested the ability of each phage to clear *Enterococcus* host cultures and prevent the emergence of phage resistant *Enterococcus*. We found that some phages which were ineffective individually were effective at clearing the bacterial culture when used in cocktails. To understand the dynamics within phage cocktails, we used qPCR to track which phages increased in abundance in each cocktail, and saw dynamics ranging from one dominant phage to even phage growth. Further, we isolated several phage-resistant mutants to test for altered Vancomycin sensitivity. We found that mutants tended to have no change or slightly increased resistance to Vancomycin. By demonstrating the efficacy of phage cocktails in suppressing growth of antibiotic susceptible and VRE clinical isolates when exposed to phages, this work will help to inform cocktail design for future phage therapy applications.

**IMPORTANCE:** Antibiotic resistant *Enterococcus* infections are a major health crisis that requires the development of alternative therapies. Phage therapy could be an alternative to antibiotics and has shown promise in *in vitro* and in early clinical studies. Phage therapy in the form of cocktails is often suggested, with similar goals as the combination therapy that has been successful in the treatment of HIV infection, but there is little understanding about how to combine phages most effectively. Here we utilized a collection of 20 *Enterococcus* phages to test whether several phage cocktails could prevent the host from evolving resistance to therapy and to determine whether evolving resistance to phages affected host susceptibility to antibiotics. We showed that cocktails of two or three unrelated phages often prevented the growth of phage-resistant mutants, when the same phages applied individually were not able to.

## INTRODUCTION

Antibiotic resistant bacterial infections have emerged as a major health crisis. Overuse of antibiotics has led to rising rates of antibiotic resistance and, therefore, steps must be taken to develop alternative therapies. The ESKAPE pathogens are six high-priority antibiotic resistant bacteria that are responsible for the majority of antibiotic resistant hospital acquired infections ^1^. Among the ESKAPE pathogens is *Enterococcus faecium*, a gram-positive bacterium responsible for several human illnesses, including sepsis, urinary tract infections (UTIs), endocarditis, wound infections and alcoholic liver disease ^2–6^. *Enterococcus* species are regular members of vertebrate gut microbiomes, and they are also pathogens of enormous clinical significance due to their high potential for resistance to antibiotics (e.g. vancomycin) ^7^. Antibiotic treatment leads to a high abundance of *Enterococcus* spp. in the gut, as many other members of a healthy gut microbial community are more sensitive to antibiotic treatment ^4,8-10^. There are multiple species of *Enterococcus* that are known to cause human disease, but the most common are *Enterococcus faecium* and *Enterococcus faecalis*. Both species are capable of developing resistance to the antibiotic vancomycin. Each of these *Enterococcus* species is known to cause disease, particularly in individuals who have been treated with multiple antibiotics. Developing alternative therapies to treat *Enterococcus* infections is of significant importance to reduce long-term infections such as endocarditis and to reduce the emergence of *Enterococcus* infections in patients who have been treated with antibiotics.

Because of their propensity for antibiotic resistance and their greater abundance in critically ill or antibiotic treated patients, *Enterococcus* is a logical focus for studies of basic phage biology. Surprisingly, there are relatively few characterized *Enterococcus* phages. Bacteriophage (phage) therapy is an alternative treatment to antibiotics that has shown promise for treating *Enterococcus in vitro* and in animal models ^11–13^. However, phage therapy is rarely used, and in many places is used only as salvage therapy, partly due to a relative paucity of basic research into its safety, mechanisms of action, and best practices of use.

Although phages are posited as a solution for antibiotic resistance, bacteria can also evolve resistance to phage infection. Bacteria exist in a constant evolutionary battle with phages, and thus have evolved many systems to resist phage infection, including preventing phage binding, restriction modification systems, CRISPR-Cas9 immunity, and abortive infection^14,15^. Given strong selective pressure from a single phage, bacteria often quickly evolve resistance to that phage in laboratory settings^16^. Phages evolve to combat and circumvent bacterial resistance mechanisms, which may provide phage therapy with an advantage over antibiotics^17^. Furthermore, the phenotypic effects of evolving resistance to phages are poorly understood. For example, phage-resistance may alter susceptibility to antibiotics^18,19^. Vancomycin resistant *Enterococcus* (VRE) that evolved resistance to phage had greatly increased susceptibility to vancomycin, in addition to mouse gut colonization defects^12^. Increasing antibiotic susceptibility would be an ideal outcome of evolving resistance to phage, but further research is required to determine how often these fitness trade-offs occur.

Phage therapy is sometimes administered as a cocktail of phages. Because many phages have narrow host ranges, multiple phage strains can be used to increase the likelihood that several or all strains of the target bacteria will be killed^20,21^. Theoretically, using a cocktail of phages could also decrease the chance that a phage-resistant mutant can arise, as multiple orthogonal resistance mechanisms would need to evolve simultaneously. Evolving a resistance mechanism can also provide a fitness disadvantage, which can make the bacteria more susceptible to cell death via other phages. Similar to phage cocktails, combinations of antibiotics are used to treat tuberculosis infections, and combinations of antivirals are used to treat HIV^22,23^. However, the efficacy of phage cocktails has not been thoroughly explored, and it is often assumed without strong evidence that a cocktail of phages will be more effective than a single phage. Thus, there are no clear design principles for crafting effective phage cocktails.

Phage therapy has shown promise in *in vitro* and *in vivo* mouse experiments^12,13,24^. Additionally, *Enterococcus* phages have been shown to be effective at disrupting *Enterococcus* in biofilms, which are generally much harder to treat than planktonic cells because antibiotics have trouble penetrating biofilms^13^. *Enterococcus* phages have also been used to treat humans. Two phage cocktails sold by the Eliava Institute of Bacteriophages, Microbiology, and Virology in Georgia were shown to contain abundant *Enterococcus* phages^25^. *In vitro, Enterococcus* phages have been shown to be more effective than single phages at preventing the growth of resistant *Enterococcus* mutants^26^. While encouraging, this represents only one example of cocktail design and does not offer any insight into cocktail design principles.

We utilized our collection of *Enterococcus* phages to test the efficacy of different combinations of phages in killing bacteria and inhibiting the proliferation of phage-resistant mutants. We tracked the abundance of each phage in a cocktail to better understand which phages may be most responsible for the cocktail’s antimicrobial activity. Finally, we measured the vancomycin susceptibility of phage-resistant mutants to determine whether evolving resistance to phage affected antibiotic susceptibility.

## RESULTS

### Host ranges of phages

We isolated phages against *Enterococcus* from influent samples from the Orange County Sanitation district, and then used plaque assays to test the phage susceptibilities of recently collected clinical isolates of *Enterococcus*, which were vancomycin resistant *Enterococcus* (VRE) or vancomycin susceptible *Enterococcus* (VSE). We identified phages that were capable of killing isolates of *E. faecium* and/or *E. faecalis*, but the *E. faecalis* isolates were generally more susceptible to a greater number of phages (**Figure 1**). As is observed often for clinical isolates, a much larger proportion of the *E. faecium* isolates were VRE compared to the *E. faecalis* isolates; however, we identified phages that were capable of killing both VRE and VSE isolates. Most phages from the *Myoviridae* family showed broad host ranges, especially Ben, Bop, and V12, which infected almost all the VRE and VSE isolates. Despite the differences between *E. faecium* and *E. faecalis* clinical isolates, we identified phages with broad host ranges that were capable of killing both species. Moreover, some of these phages, from *Siphoviridae, Myoviridae*, and *Podoviridae* families, were capable of killing highly antibiotic resistant VRE isolates.

**Figure 1.**
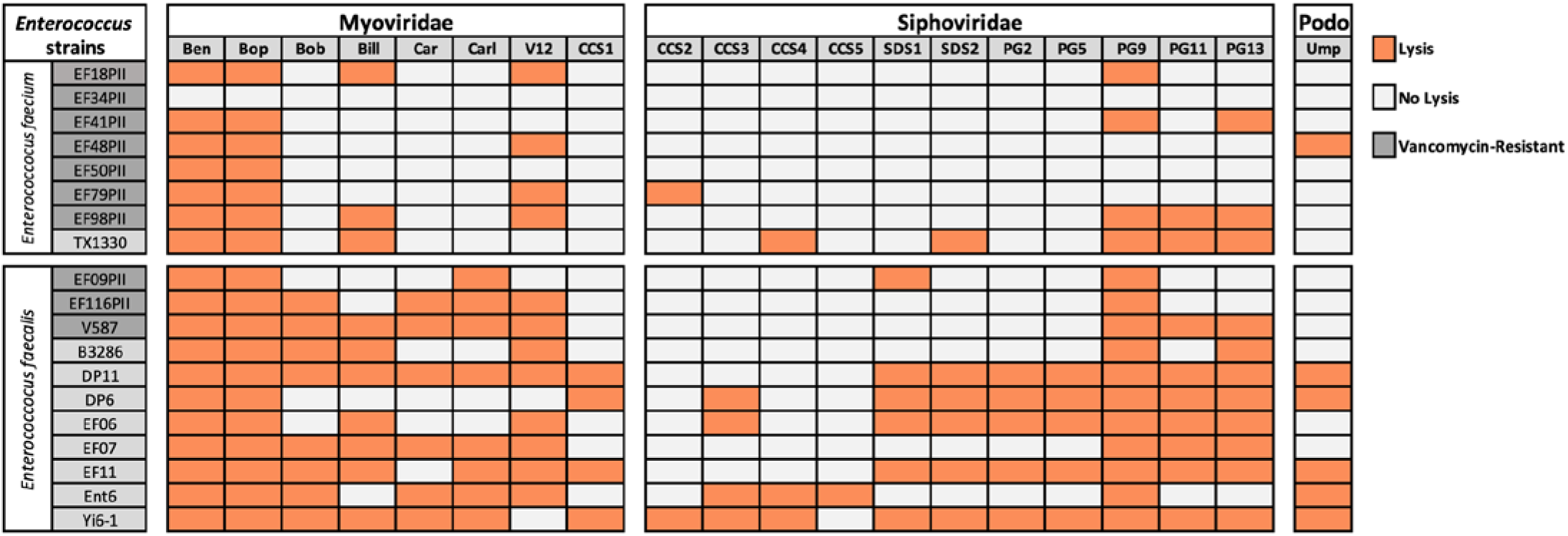
Host ranges of *Enterococcus* phages as determined using plaque assays. Complete lysis is indicated by orange boxes and white boxes represent no lysis. Several vancomycin sensitive (light grey boxes around strain names) and vancomycin resistant (dark grey boxes around strain names) *Enterococcus* isolates were used for this study.

### Phage cocktails reduce the evolution of resistant mutants

To understand the advantage of using phage cocktails over single phages, we measured bacterial growth over 72 h in liquid media to observe phage lysis and detect the emergence of resistance (as seen by bacterial growth). Four *E. faecalis* strains were chosen for testing the efficacy of phage cocktails, with strain Yi6-1 allowing for the most combinations because it was susceptible to 18 of the 20 phages in our collection (**Figure 1**). The multiplicity of infection (MOI) of phage cocktails tested on *E. faecalis* strains, including 0.1, 0.01, and 0.001, did not consistently affect the emergence of phage-resistant mutants (**Supplemental Figure S1**). Therefore, we chose to infect strains at the highest MOI of 0.1. When susceptible *Enterococcus* was infected with a single phage at MOI 0.1 in liquid media, its growth is initially stunted compared to the no-phage control, but if resistance emerges, growth is often observed within 24 to 60 hours (data not shown). In a minority of instances, such as was observed for phages Bop, CCS3, SDS2, and Car, a single phage was able to prevent measurable host growth over the 72 hour period. However, for most other instances, the growth of phage-resistant *Enterococcus* emerged within 72 hours of exposure to a single phage (**Figure 2**). There were no single phages that were able to prevent host growth over the entire 72 h period in all four *E. faecalis* strains tested.

**Figure 2.**
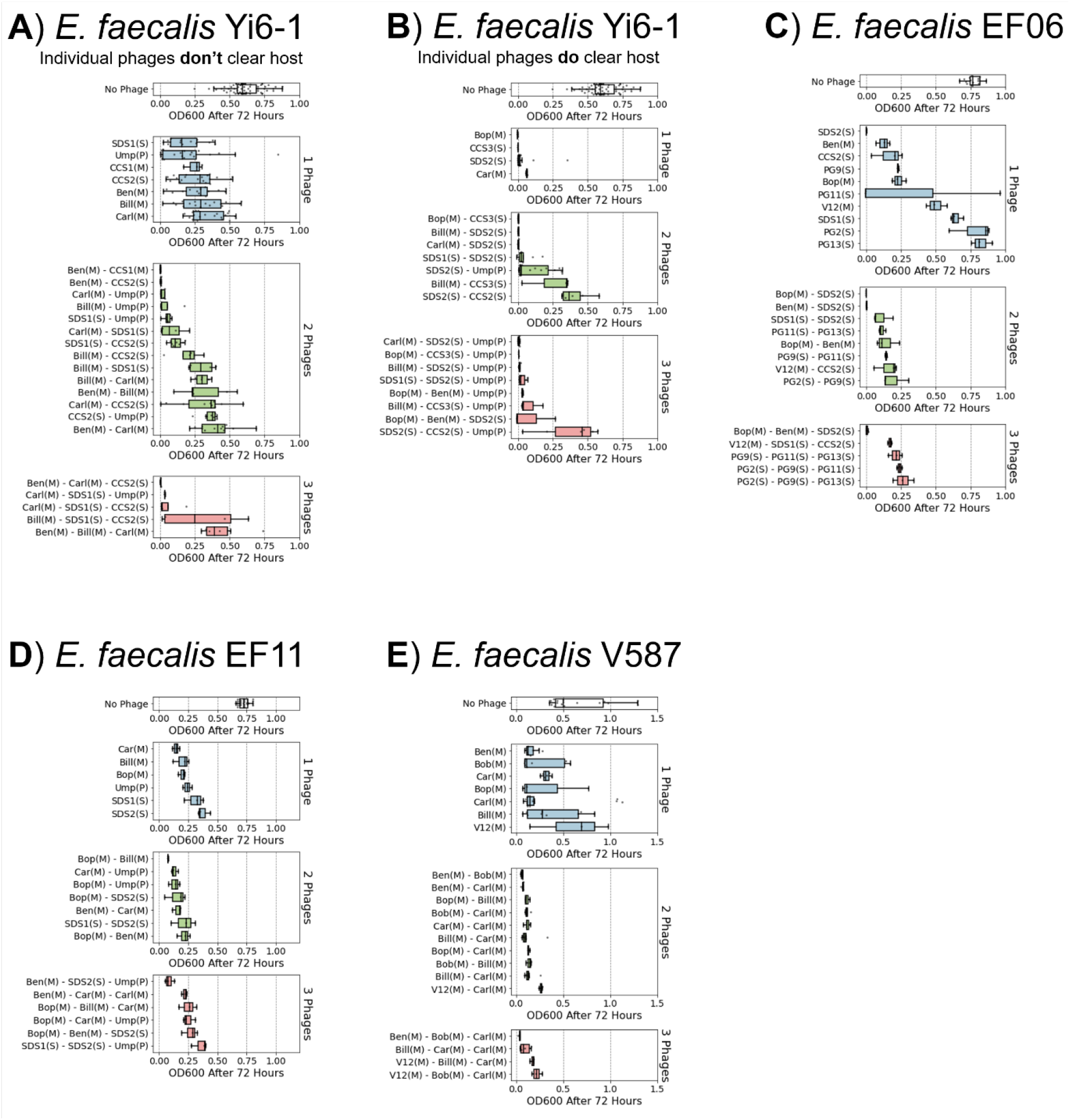
Phage cocktails clear cultures and prevent growth of *E. faecalis* for 72 h. Combinations of one, two, or three phages were added to susceptible *E. faecalis* hosts in exponential growth phase and bacterial growth after 72 h was measured by OD_600_. **A**) *E. faecalis* Yi6-1 cultures in which the individual phages are unable to prevent growth. **B)** *E. faecalis* Yi6-1 cultures in which the individual phages are able to prevent growth. **C**) *E. faecalis* EF06. **D**) *E. faecalis* EF11. **E**) *E. faecalis* V587 (Vancomycin resistant strain/VRE). (M): *Myoviridae* phage, (S): *Siphoviridae* phage, (P): *Podoviridae* phage.

When *E. faecalis* Yi6-1 was infected with combinations of two phages, some combinations consistently prevented the growth of phage-resistant mutants, while other combinations failed to do so (**Figure 2A-B**). For *E. faecalis* strains Yi6-1, EF06, and EF11, successful two-phage cocktails were usually composed of multiple phage families (for instance, *Myoviridae* and *Siphoviridae*), but not all cocktails composed of two different families were successful at preventing growth (**Figure 2A-D**). *E. faecalis* strain V587 (VRE) was only tested with *Myoviridae* phages, as it was not susceptible to any of the *Siphoviridae* or *Podoviridae* phages in our collection. In V587, some single-family combinations of *Myoviridae* phages were effective at preventing growth (**Figure 2E**). Three-phage cocktails were often successful at preventing bacterial growth for all four *E. feacalis* strains (**Figure 2**). However, adding a third phage to the cocktail did not generally improve a cocktail’s ability to prevent the emergence of phage resistance, as most successful three-phage cocktails contained a phage combination that worked together as two-phage cocktails.

### *Siphoviridae and Podoviridae* phages dominate in most cocktails

To understand the dynamics of our cocktails, we used quantitative PCR (qPCR) to estimate the relative abundance of each phage in the cocktail during the course of infecting *E. faecalis* strain Yi6-1. We selected cocktails containing two or three phages that were ineffective as individual phages, but were successful as cocktails in preventing bacterial growth over 72 hours. Phage DNA was extracted immediately after phage and host inoculation (0 hours), as well as at 24, 48, and 72 hours post-inoculation. Changes in phage abundances over time in each of eight cocktails were plotted over 72 hours, as compared with the phage-only control (**Figure 3**). Optical density (OD) measurements for each culture were also plotted to demonstrate that each phage cocktail largely prevented the growth of Yi6-1 for the duration of 72 hour period.

**Figure 3.**
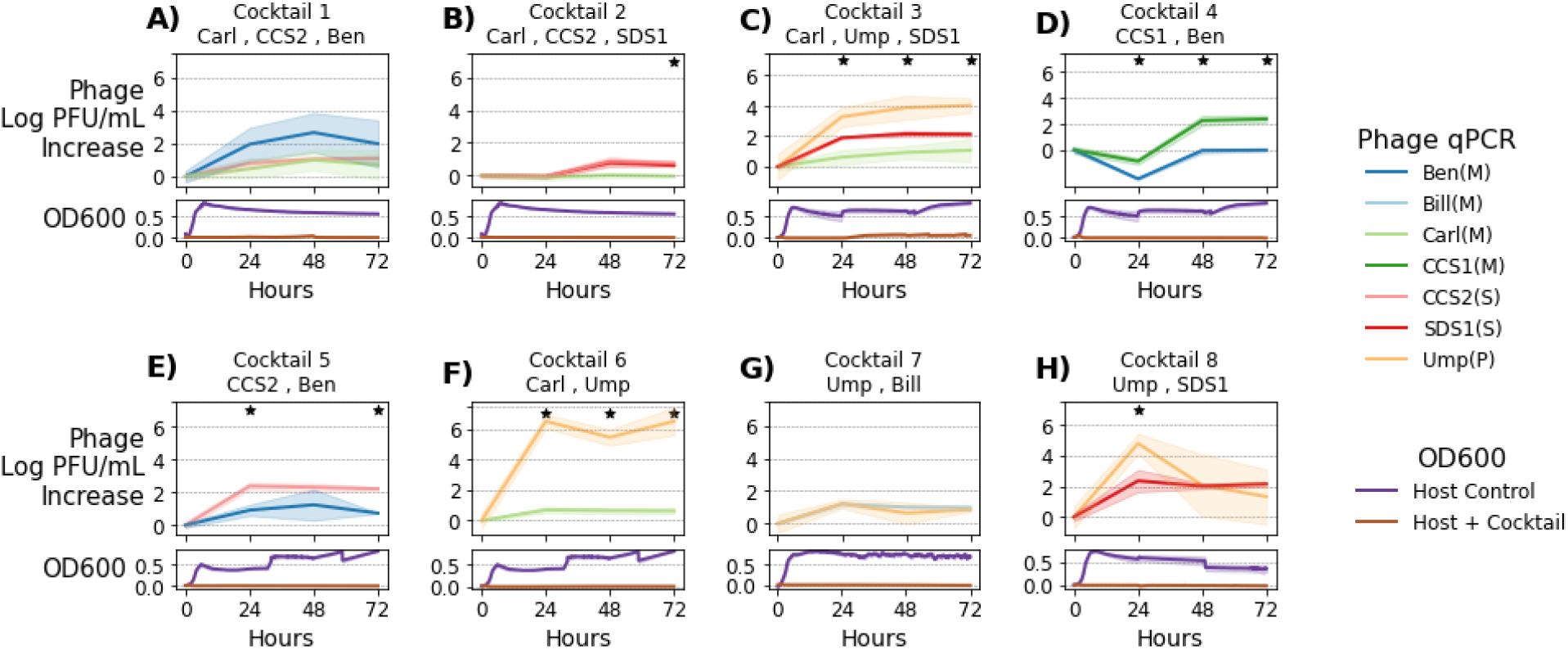
Tracking phage concentration changes within phage cocktails. Eight different cocktails of two or three phages were added to *E. faecalis* Yi6-1 cultures and the concentration of each individual phage was measured by qPCR every 24 h for 72 h of growth. Upper panels represent change in phage concentration and lower panels represent host growth as measured using OD_600_ at 0, 24, 48, and 72 h post-inoculation. Solid lines represent the mean (n=3) increase in each phage’s abundance since hour 0. Lighter color ribbons show standard deviation. Stars indicate significant differences between individual phage increases as determined using ANOVA (p < 0.05).

Cocktail 1 contained phages Carl (*Myoviridae*), CCS2 (*Siphoviridae*), and Ben (*Myoviridae*). While the concentrations of Carl and CCS2 increased by ~1.5 log PFU/ml, the concentration of Ben increased by ~3 log PFU/ml (**Figure 3A**), suggesting that it was the most active phage in this cocktail. In cocktail 2 with phages Carl (*Myoviridae*), CCS2 (*Siphoviridae*), and SDS1 (*Siphoviridae*), change in phage concentrations remained at or below 0 PFU/ml for the first 24 h, despite preventing any detectable bacterial growth. However, at 48 and 72 h, phages CCS2 and SDS1, but not Carl, had increased abundances (**Figure 3B**). In the third phage cocktail with phages Carl (*Myoviridae*), Ump (*Podoviridae*), and SDS1 (*Siphoviridae*), all three phages increased in concentration over time, with Ump increasing the most (~4 log PFU/mL), SDS1 in the middle (~2 log PFU/mL), and Carl increasing the least (~1 log PFU/mL) (**Figure 3C**). Cocktail 3 was the only cocktail in which any bacterial growth was observed, starting around 24 h in two of the three replicates and remaining at a low OD_600_ for the remainder of the experiment, significantly lower than the OD_600_ of >0.5 often observed when resistance emerged.

We also monitored the concentration of each phage in successful two-phage cocktails. In cocktail 4, *Myoviridae* phages CCS1 and Ben increased by similar amounts, with CCS1 having a slight edge (**Figure 3D**). In cocktail 5, *Siphoviridae* phage CCS2 consistently increased by ~1 log PFU/mL more than phage Ben (**Figure 3E**). Our data suggested that the phages like CCS1 and CCS2 consistently and strongly reduced bacterial growth, as their concentrations increased significantly (from 2 to ~4 and 0 to ~2.5 log PFU/ml, respectively) compared to Ben (only increased from 1.5 to ~2 log PFU/ml) (**Figure 3D, E**). In other two-phage cocktails 6 (Carl-Ump), 7 (Bill-Ump), and 8 (SDS1-Ump), the *Podoviridae* phage Ump was found to have greater increases in its relative abundance (the concentration increased from 1.5 to ~ 6 log PFU/ml) compared to *Myoviridae* phages Carl and Bill (remained stable at ~2 log PFU/ml throughout the duration of infection). However, for the SDS1-Ump cocktail, a slight increase in the concentration of *Siphoviridae* phage SDS1 was observed, though still less than the change in concentration of *Podoviridae* phage Ump.

### Resistance to phage can modulate antibiotic susceptibility

We examined six *Enterococcus* phage-resistant mutants generated from three wild-type *Enterococcus* strains in the cocktail experiments to determine whether some may have altered susceptibility to vancomycin. *E. faecalis* strains DP11 and Yi6 were initially susceptible to vancomycin while *E. faecalis* V587 was initially vancomycin resistant. First, mutants were sequenced to determine the mechanism of increased phage resistance (**Table 1**). Five out of six resistant mutants had mutations in the Epa exopolysaccharide synthesis locus. Phage-resistant *Enterococcus* mutants often had similar or slightly decreased levels of susceptibility to vancomycin (**Figure 4**). Many phage-resistant *Enterococcus* mutants did not achieve the same growth characteristics (OD_600_ compared to the wild-type isolate), indicating that these mutants had growth defects.

**Figure 4.**
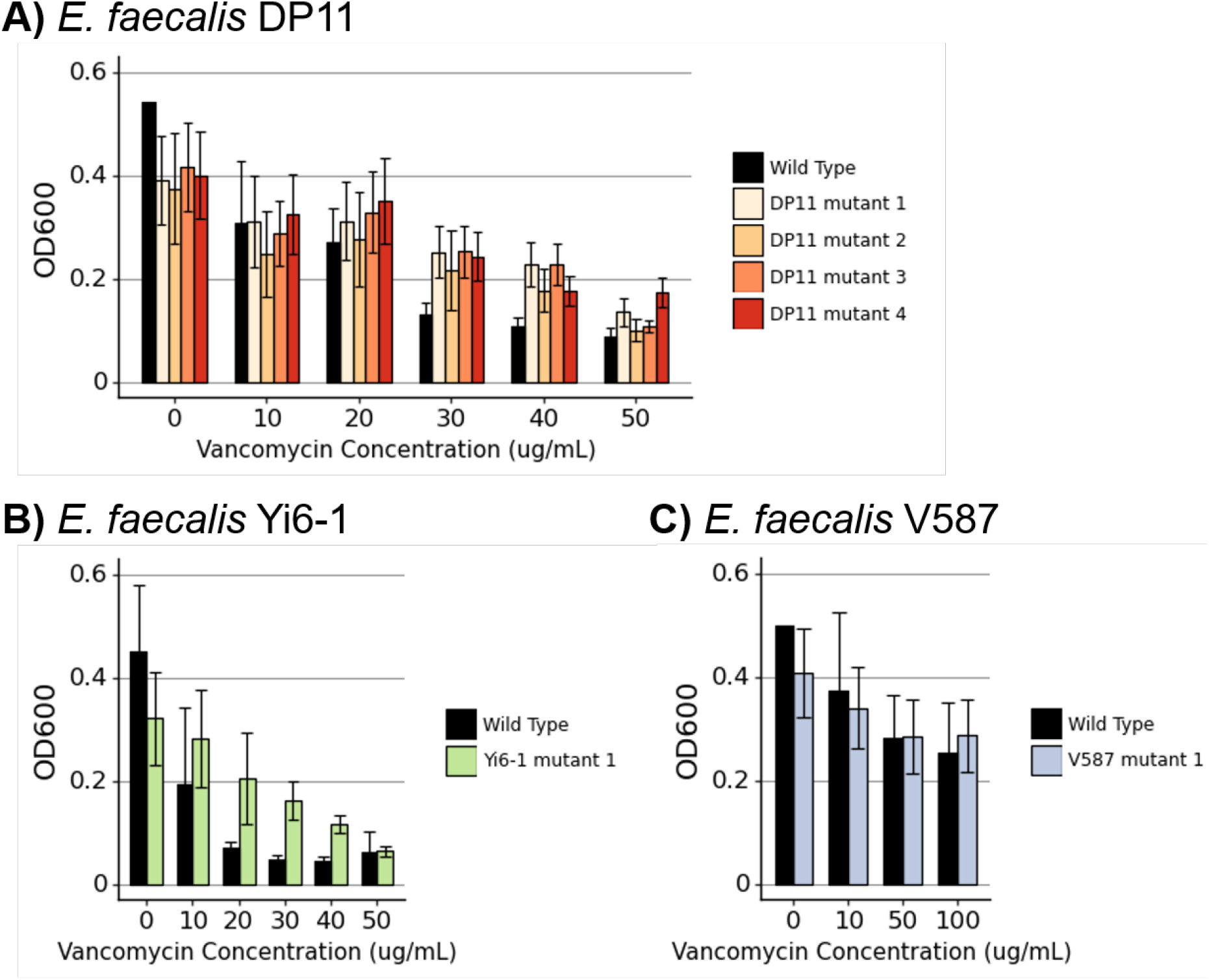
Vancomycin susceptibility of *Enterococcus* mutants. Bacterial growth with increasing concentrations of Vancomycin of phage-resistant *Enterococcus* mutants generated from **A)** *E. faecalis* DP11, **B)** *E. faecalis* Yi6-1, and **C)** vancomycin resistant strain *E. faecalis* V587. Bars represent means and error bars represent standard deviations of three biological replicates.

**Table 1.**
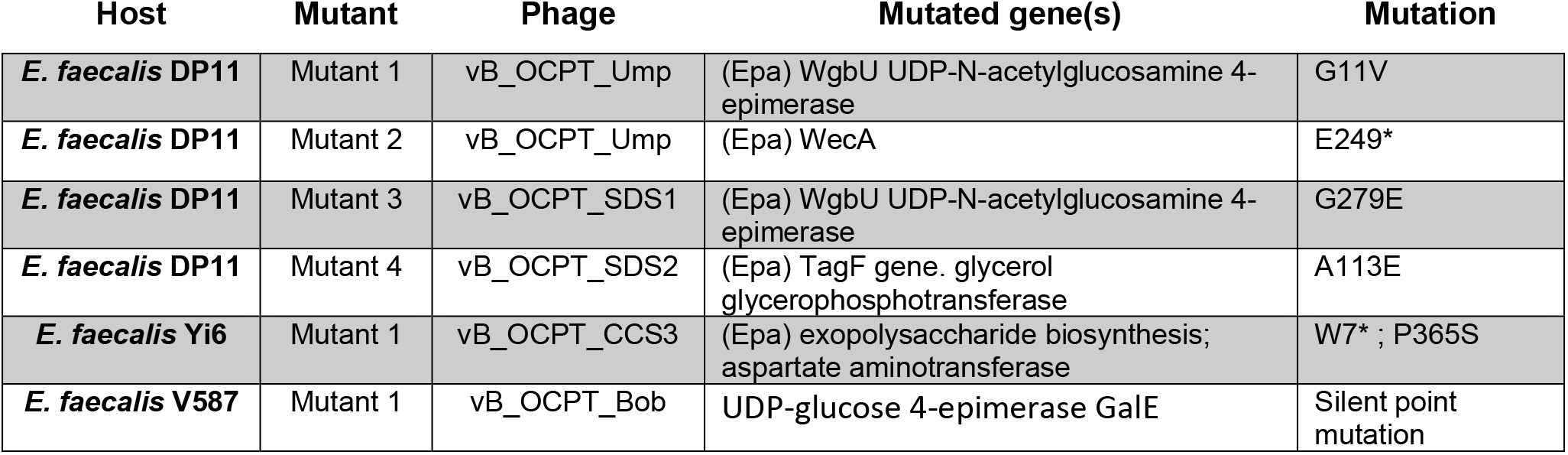
Mutations in *Enterococcus* providing phage resistance. Genes that are part of the Epa exopolysaccharide synthesis locus are denoted with (Epa).

## DISCUSSION

Some phage cocktails have previously been shown to be effective at killing bacteria and preventing the growth of resistant mutants, but little focus has been placed on the design principles for effective cocktails. Here, we showed that phage cocktail design is an important consideration because not all combinations of phages consistently prevent the growth of phage-resistant mutants.

While using phage cocktails including a diversity of phages targeting multiple disparate bacterial types has been a common practice, there is no standard for how many phages should be used. The pyophage (PYO) cocktail from the Georgian Eliava Institute of Bacteriophages has been shown to contain approximately thirty different phages targeting multiple different bacterial hosts^25^. Recent uses of phage therapy designed to target a single strain of a bacterial species generally include between one and six phages ^27–29^. Often there is no obvious rationale behind the number of phages chosen to administer during phage therapy, although lack of access to phage that can infect a host of interest may be an important limitation. Here, we show that combinations of two phages are often enough to prevent the growth of phage-resistant mutants. Using more than two diverse phages in a cocktail would increase the chances of choosing two phages that displayed synergy in preventing the growth of phage-resistant mutants. However, increasing the number of phages could pose the risk of antagonistic interactions between phages^30^.

Several approaches exist for optimizing phage cocktail design. Experimental evolution of a phage can result in mutant phages with expanded host ranges, which can be used in phage cocktails^31,32^. Another approach is to use phage-resistant hosts to isolate new phages with complementary host ranges^33^. Synthetic approaches can also be effective, such as using site-directed mutagenesis to create phages that bind to different sites and would thus complement each other in a phage cocktail^34^. Our experiments relied on the diversity of our phage collection to create cocktails, but many bacterial hosts lack large collections of characterized phages. Evolutionary or synthetic approaches would be useful for expanding phage catalogs for specific hosts.

Some phages in our collection were effective against *E. faecalis* and *E. faecium*, including both VRE and VSE isolates. This suggests that these phages would be good candidates for phage cocktails to eradicate a variety of *Enterococcus* isolates. If the phages in a cocktail attach to different bacterial target proteins, a cocktail of diverse phages is more capable of lysing the pathogen even when a mutant arises. As seen in **Figure 2A**, combinations of phages that individually cannot clear a culture of *E. faecalis* Yi6-1 are able to do so when used in combinations. This evidence shows that the evolutionary advantage of phage-resistant mutants can be diminished with the use of well-designed cocktails. While this is a well-known phenomenon that exists for antibiotics in the treatment of diseases such as tuberculosis, our *in vitro* evidence suggests that it might also be applicable to phage therapy.

Comparing the abundances of each phage over time in the eight phage cocktails shown in **Figure 3** yields a wide range of outcomes, from one phage greatly outpacing the other (cocktail 6), to relatively even abundances (cocktails 1,7, 8). All the phages used in these cocktails were ineffective at clearing *Enterococcus* cultures alone, yet all of these cocktails, with their varied phage dynamics, resulted in clearing the *Enterococcus* culture. This shows that there is not one single path to an effective phage cocktail. Even if one phage appears to be dominating, the other phage is still necessary to prevent the emergence of a phage-resistant mutant.

Phages that display synergy with antibiotics might be ideal for phage therapy. Synergy can occur if the evolution of resistance to phage comes ast the cost of increased susceptibility to antibiotics, or vice versa. The most famous example of this is a *Pseudomonas aeruginosa* phage that binds to an antibiotic efflux pump, resulting in a fitness trade off where bacteria can evolve resistance to phage by losing the antibiotic efflux pump^18^. Several studies, including this one, have shown *Enterococcus* evolving resistance to phage infection through mutations in the Epa locus. While not observed in this study, Epa mutations have been seen to increase susceptibility to vancomycin in resistant strains^12,35-37^. The mechanism by which this fitness trade-off occurs is not clear; vancomycin targets peptidoglycan and the Epa locus regulates exopolysaccharide synthesis, so both interactions occur at the cell wall^38–40^. In this study, evolution of phage resistance in *Enterococcus* was not seen to sensitize VRE, and slightly increased vancomycin resistance in VSE. Different Epa mutations may have different effects on antibiotic susceptibility. In addition, the genetic background of the *Enterococcus* strain likely affects the phenotypic outcomes of the evolution of phage resistance. Synergistic outcomes of phage-host co-evolution that result in better treatment options are far from guaranteed, but the enormous numbers of understudied phage-host interactions leave room for the discovery of phages that co-evolve with their pathogenic bacterial hosts in ways that make infection treatment more tractable.

The need for alternative therapies for antibiotic resistant bacterial infections has never been greater than it is right now with the crisis of antibiotic resistance upon us. Bacteriophages represent an alternative with the potential to overcome some of the shortcomings of antibiotics, but there are significant knowledge gaps that reduce the utility of phages in humans. Problems that face phage therapy in the future include the emergence of host resistance, limited information on how phages may collectively eradicate their hosts, and a lack of guidance on how many phages may be necessary in cocktails to achieve phage therapy goals. Using the model of the diverse bacterium *Enterococcus*, which has multiple species capable of causing debilitating human infections, and has the ability to acquire significant antibiotic resistance, we have demonstrated that phages have significant potential to kill their hosts and reduce the emergence of resistant isolates. For *Enterococcus*, our data suggest that host killing was not substantially increased by adding more than two phages to cocktails, but a larger study is needed to confirm these findings. Rationally designed phage cocktails hold the potential to significantly advance phages as antibiotic alternatives for the treatment of human pathogens such as *Enterococcus*.

## MATERIALS AND METHODS

### Bacterial strains and culture conditions

The bacterial strains used in this study are listed in **Supplemental Table S2**. Most of the *E. faecalis* and *E. faecium* strains were identified from plates that were sent to the UCSD Center for Advanced Laboratory Medicine for identification by MALDI-TOF, where the reported antibiotic susceptibilities were also assessed. A few of the strains were obtained from the Human Microbiome Project repository at the Biodefense and Emerging Infections Research Resources Repository (BEI: www.beiresources.org). *E. faecalis and E. faecium* strains were cultured in Brain Heart Infusion (BHI) broth. All strains were grown at 37°C in liquid medium overnight with shaking. Solid medium was prepared with 1.5% agar when culturing bacteria or 1.0% bottom agar and 0.3% top agar for plaque assays.

### Host Range Determination

Phage susceptibilities were measured in several clinically relevant bacterial strains (**Supplemental Table S2**) that demonstrated multi-drug resistance, causing illness. Phage susceptibility was determined using a spot assay in which 5 μl of each phage lysate was spotted on a lawn of an *Enterococcus* strain on a 1.5% Agar plate infused with BHI. The spots were allowed to dry at room temperature for 30 min before being incubated at 37 °C for 24 h. The next day, plates were examined to identify the host’s susceptibility based on the appearance of the zones of cell lysis.

### Determination of Multiplicity of Infection (MOI)

Multiplicity of infection is defined as the ratio of the number of phage particles to the number of target cells of each host. We tested three MOI (0.1, 0.01 and 0.001) as follows. A single colony was picked from a streak plate and was grown overnight in BHI broth at 37 °C. The next day, the bacterial culture was diluted to 0.05 OD_600_ in fresh BHI and once the initial dilution was completed, the serial dilution of 10^-1^ to 10^-8^ of each bacterial culture was performed using Phosphate Buffered Saline. For each isolate, the aliquot from serial dilutions of 10^-4^ and 10^-5^ were plated on 1.5% BHI agar plates and were incubated overnight at 37 °C. The next day, the number of colonies were counted on each dilution plate and the number colony forming units (CFU/ml) was determined (**Supplemental Table S1**). A plaque assay was then performed for all the phages with their respective isolates to determine host susceptibility to phages. The cells from the log phase were infected with different phages at different dilutions. Based on plaque assay plates, we determined the titer of each phage (PFU/ml) (**Supplemental Table S1**). Based on observed colony forming units (CFU/ml) of *Enterococcus* isolates and the titer of each phage (PFU/ml), the appropriate volume of isolates and phages were determined at three different MOI: 0.1,0.01 and 0.001 to carry out growth curve experiments.

### Phage Cocktails

Cocktails consisting of one, two, or three phages were tested against *E. faecalis* Yi6-1, EF06, EF11 and V587. A single colony was inoculated into BHI broth and grown overnight in a shaker at 37 °C. The next day, bacterial cultures were diluted to 0.05 OD_600_ in fresh BHI. On the basis of three different MOIs, appropriate volumes of *Enterococcus* and phages were determined and combined in a single well inside a 96-well plate along with enough BHI to make up a total volume of 200 μl. For two-phage cocktails, half of previously determined volumes of each of the two unique phage stocks were added. To conduct three-phage cocktails, one third of previously determined volumes of each of the three unique phage stocks were added. To avoid desiccation in edge wells of the 96 well plates used for growth curve experiments, 200 μl of fresh BHI media was added into these edge wells to avoid desiccation issues in wells adjacent to edge wells.

### Quantitative PCR (qPCR)

Phage cocktails were grown in BHI for 72 h with their respective hosts (Yi6-1, EF06 and EF11) and this experiment was carried out in 96-well plates with three biological replicates. In the positive control, phages were added without a host. For the negative control, appropriate hosts were grown in BHI for 72 h in the absence of phage. During this experiment, the respective samples were collected at four time points: 0 h, 24 h, 48 h, and 72 h. From the samples collected at each time point, total genomic DNA was isolated using the DNeasy Blood and Tissue Kit (Qiagen). For the unknown standards, we performed a plaque assay for all the phages to find their concentrations in PFU/ml and the genomic DNA of the standards were isolated using the same kit. Standard curves were generated with serial dilutions of phage (10^-1^ - 10^-8^). Phage specific primers were designed using Geneious software and are listed in **Supplemental Table S4**. The qPCR experiment was performed in 96-well PCR plate using Eppendorf Mastercycler® RealPlex with SYBR Green PCR Master Mix (Eppendorf, USA) as per the universal SYBR Green qPCR protocol where fluorescent product is detected during the last step of each cycle. The obtained melting curve data were analyzed using Eppendorf Mastercycler® RealPlex to calculate the Ct values. Ct values of the standards were then used to generate standard curves correlating log PFU/ml to Ct values, which was used to estimate the concentration of each phage in the cocktails.

### Generation of phage resistant *Enterococcus* mutants in Liquid Cultures

*Enterococcus* colonies were grown overnight and diluted to optical density at 600 nm (OD_600_) of 0.05. We added 190 μl of *Enterococcus* and 10 μl of a highly concentrated individual phage stock into each well. 96-well plates co-inoculated with bacteria and phage/two-phage cocktail were then incubated at 37°C in a spectrophotometer, where cell density, as measured by an absorbance at OD_600_, was taken every 10 min for 24-72 h. During the 1-3-day incubation, any *Enterococcus* sp. culture that showed growth after a period of no growth was streaked onto 1.5% BHI-agar plates. Colonies that grew on these streak plates were considered phage resistant mutants and were used in subsequent mutant assays.

### Vancomycin Susceptibility

To determine whether phage resistance mutations affected Vancomycin susceptibility or resistance, we cultured wild-type and mutant *Enterococcus* across a spectrum of relevant vancomycin concentrations. Wild-type and mutant *Enterococcus* colonies were grown overnight with BHI broth in a shaker at 37°C. The next day, all cultures were diluted to 0.05 OD_600_ in fresh BHI. In a 96 well plate, 190 μl of *Enterococcus* and 10 μl of Vancomycin in a gradient from 0.2 to 200 μg/ml was added in triplicate to wells and incubated in a spectrophotometer at 37°C for 10 h.

### Enterococcus Mutant Sequencing

DNA was extracted from *Enterococcus* mutants using the Quick-DNA Microprep Kit (Zymo #D3020). Before *Enterococcus* DNA extraction, lysozyme was added to lysis buffer at a concentration of 100 μg/ml and incubated at 37°C for 30 min. Libraries were prepared using a scaled-down protocol with the Illumina Nextera enzyme^41^. Short read length (75 bp) paired-end sequencing was performed on the Illumina NextSeq using the Mid Output v2 reagents. Approximately 1 million reads were obtained per sample, resulting in about 10-fold coverage across the *Enterococcus* genome.

### Sequencing Analysis

DNA sequencing reads from each phage-resistant host were aligned to their wild-type genome using Bowtie2^42^. Variants were called and filtered with samtools and bcftools and manually inspected using Geneious^43–45^. All mutations are reported in **Table 1**.

## DATA AVAILABILITY

Data from bacterial growth assays, phage qPCR, and code for analysis and making figures are available at https://github.com/swandro/phage_cocktails. Genomes for bacterial and phage strains used in this study will be uploaded to Genbank (Table S1, S2).

## ACKNOWLEDGEMENTS

We would like to acknowledge the Orange County Sanitation District for providing influent samples from which some of the phages were isolated, a T32 training grant to Stephen Wandro (T32AI007319), an R21 awarded to Katrine Whiteson and David Pride (R21AI149354) and the UC San Diego Health Clinical Microbiology Laboratory.

---

**Supplemental Figure S1.**
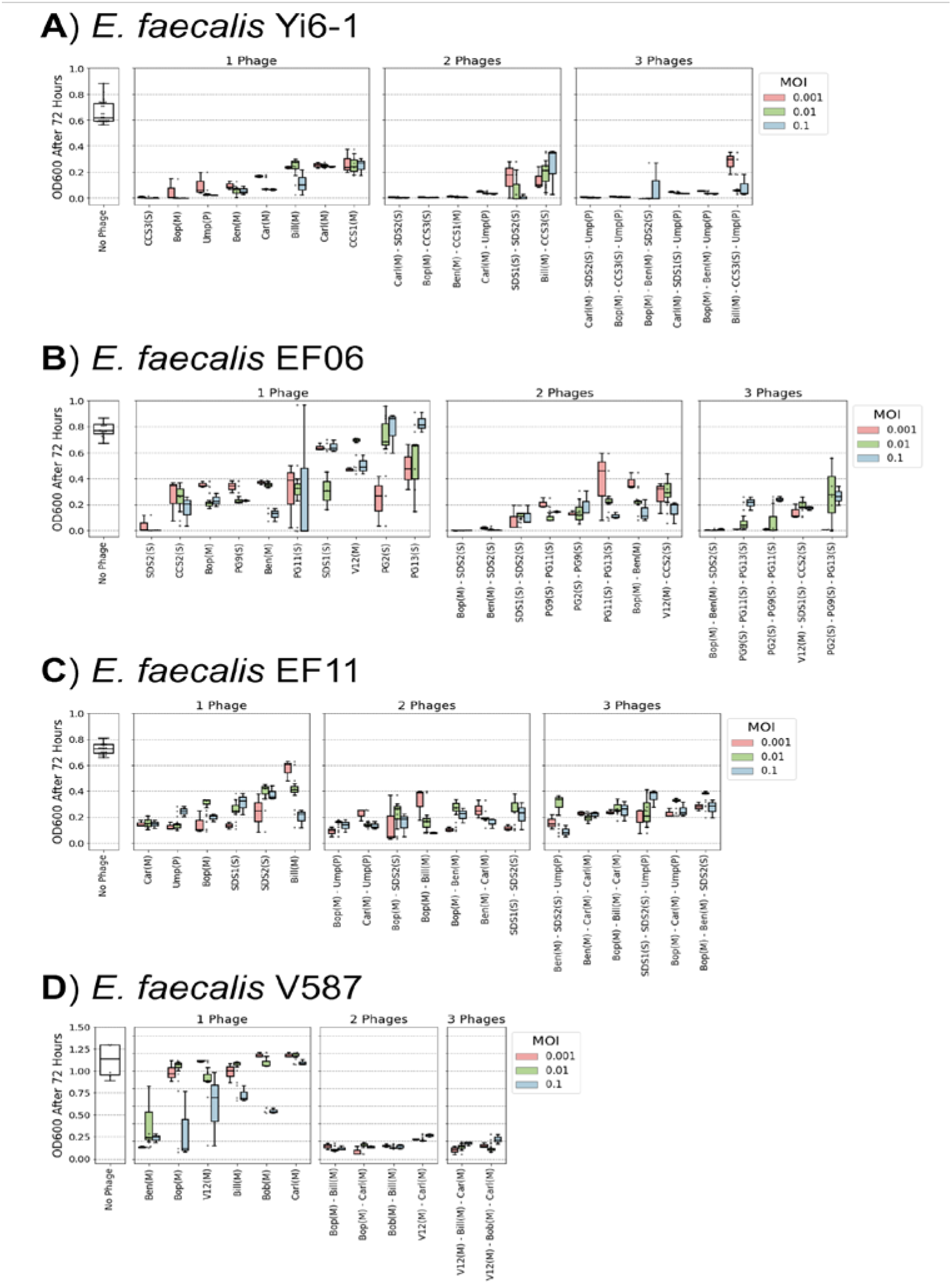
Effect of MOI on bacterial growth in phage cocktails. Three different MOIs (0.1, 0.01, 0.001) were used to study the efficiency of single phages and phage cocktails against four different *E. faecalis* isolates over the 72 h incubation period: **A)** Yi6-1, **B)** EF06, **C)** EF11, and **D)** V587.

**Supplemental Table S1.**
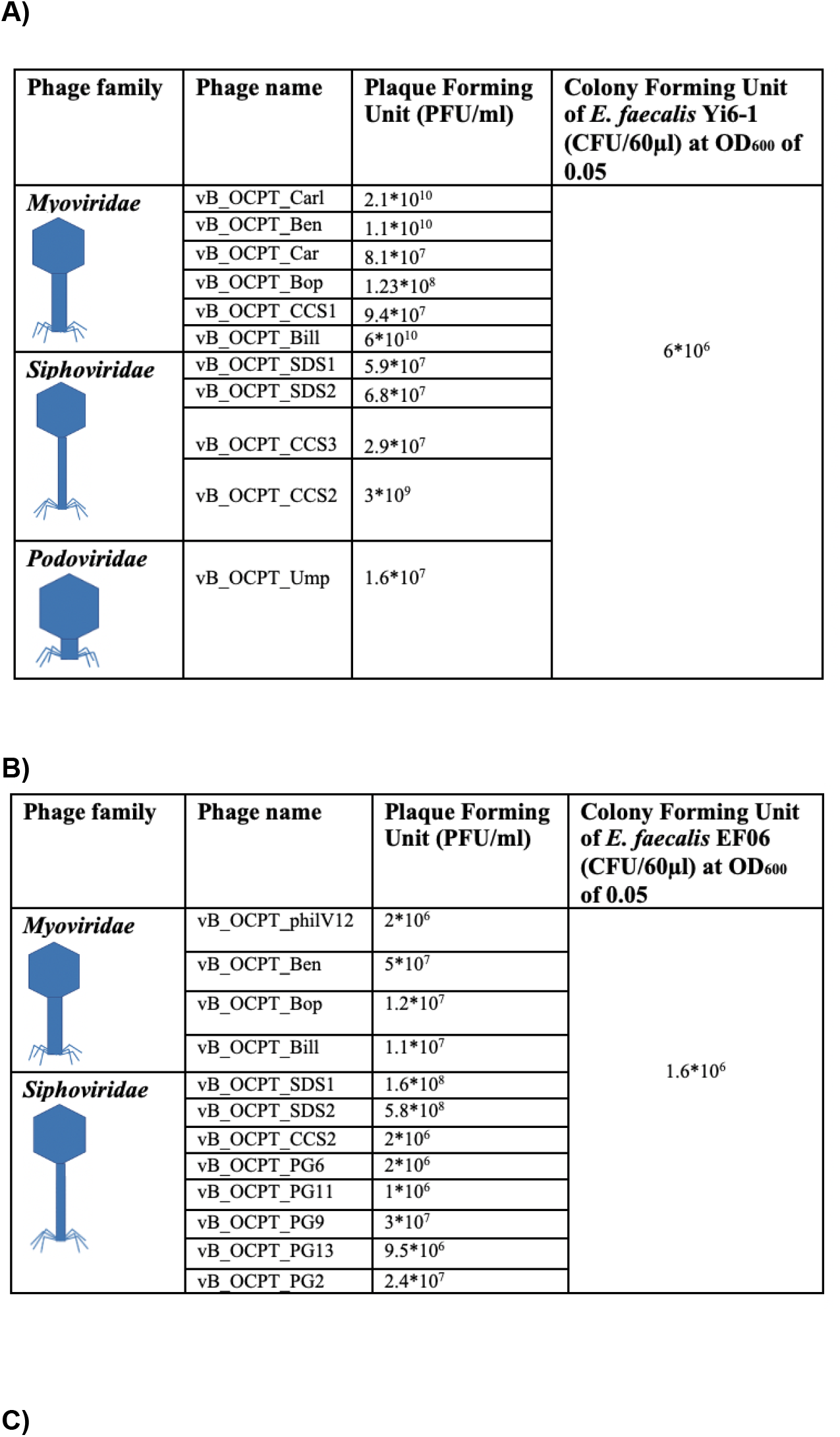
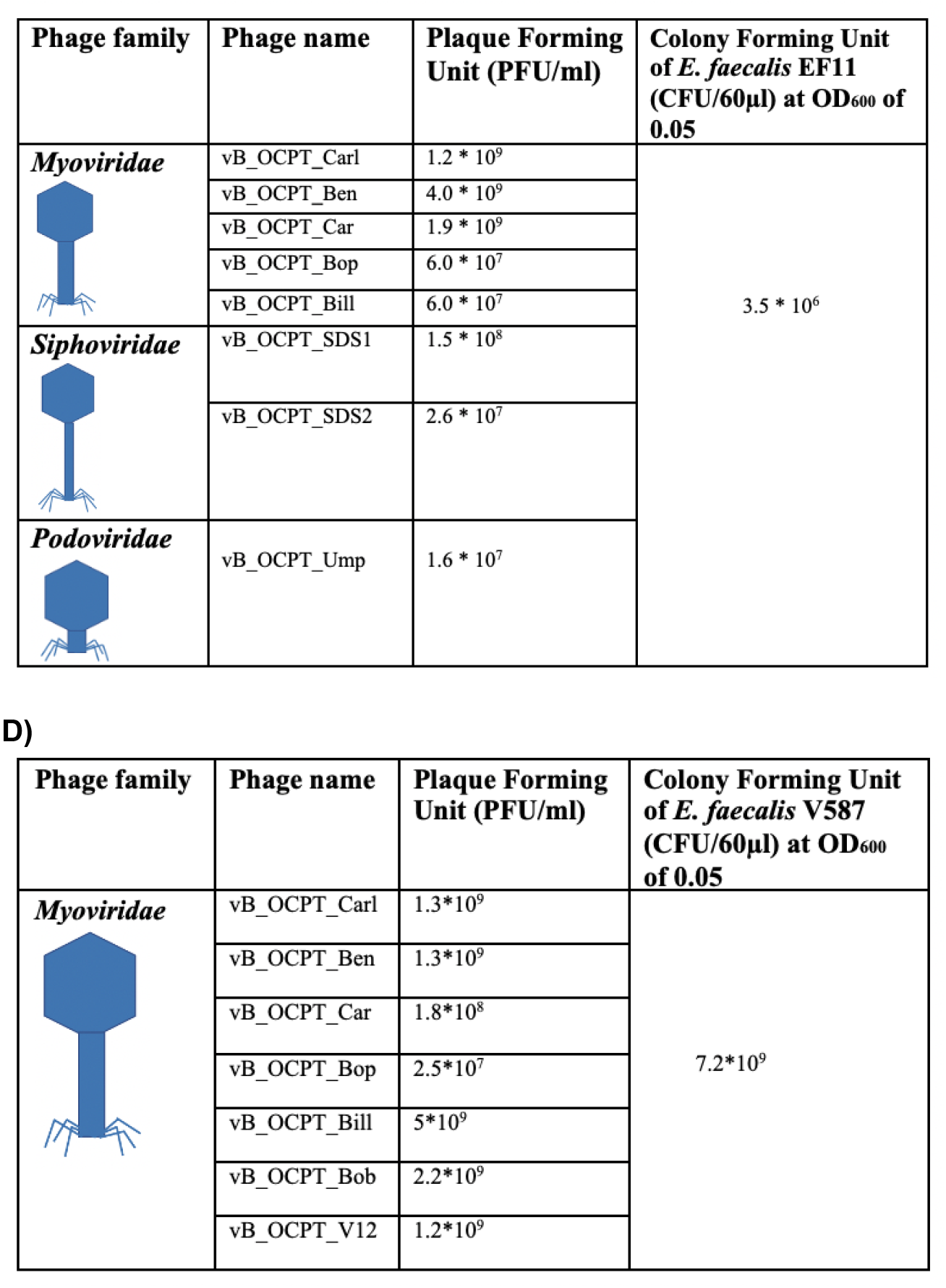
Phage titers and the concentration of the host. The phage titers (Plaque forming units (PFU)/ml) were determined using plaque assays. Viable counts of the hosts were measured by counting colonies formed from liquid cultures at an OD_600_ of 0.05 (CFU/60ul). All phages were titered on their corresponding susceptible *E. faecalis* hosts: **A)** Yi6-1, **B)** EF06, **C)** EF11, and **D)** V587.

| Phage family | Phage name | Plaque Forming Unit (PFU/ml) | Colony Forming Unit of *E. faecalis* Yi6-1 (CFU/60μl) at OD<sub>600</sub> of 0.05 |
| --- | --- | --- | --- |
| **Myoviridae** | vB_OCPT_Carl | 2.1*10<sup>10</sup> | 6*10<sup>6</sup> |
|  | vB_OCPT_Ben | 1.1*10<sup>10</sup> |  |
|  | vB_OCPT_Car | 8.1*10<sup>7</sup> |  |
|  | vB_OCPT_Bop | 1.23*10<sup>8</sup> |  |
|  | vB_OCPT_CCS1 | 9.4*10<sup>7</sup> |  |
|  | vB_OCPT_Bill | 6*10<sup>10</sup> |  |
| **Siphoviridae** | vB_OCPT_SDS1 | 5.9*10<sup>7</sup> |  |
|  | vB_OCPT_SDS2 | 6.8*10<sup>7</sup> |  |
|  | vB_OCPT_CCS3 | 2.9*10<sup>7</sup> |  |
|  | vB_OCPT_CCS2 | 3*10<sup>9</sup> |  |
| **Podoviridae** | vB_OCPT_Ump | 1.6*10<sup>7</sup> |  |

**B)**
| Phage family | Phage name | Plaque Forming Unit (PFU/ml) | Colony Forming Unit of *E. faecalis* EF06 (CFU/60μl) at OD<sub>600</sub> of 0.05 |
| --- | --- | --- | --- |
| **Myoviridae** | vB_OCPT_philV12 | 2*10<sup>6</sup> | 1.6*10<sup>6</sup> |
|  | vB_OCPT_Ben | 5*10<sup>7</sup> |  |
|  | vB_OCPT_Bop | 1.2*10<sup>7</sup> |  |
|  | vB_OCPT_Bill | 1.1*10<sup>7</sup> |  |
| **Siphoviridae** | vB_OCPT_SDS1 | 1.6*10<sup>8</sup> |  |
|  | vB_OCPT_SDS2 | 5.8*10<sup>8</sup> |  |
|  | vB_OCPT_CCS2 | 2*10<sup>6</sup> |  |
|  | vB_OCPT_PG6 | 2*10<sup>6</sup> |  |
|  | vB_OCPT_PG11 | 1*10<sup>6</sup> |  |
|  | vB_OCPT_PG9 | 3*10<sup>7</sup> |  |
|  | vB_OCPT_PG13 | 9.5*10<sup>6</sup> |  |
|  | vB_OCPT_PG2 | 2.4*10<sup>7</sup> |  |

**C)**
| Phage family | Phage name | Plaque Forming Unit (PFU/ml) | Colony Forming Unit of *E. faecalis* EF11 (CFU/60µl) at OD<sub>600</sub> of 0.05 |
| --- | --- | --- | --- |
| ***Myoviridae*** | vB_OCPT_Carl | 1.2 * 10<sup>9</sup> | 3.5 * 10<sup>6</sup> |
|  | vB_OCPT_Ben | 4.0 * 10<sup>9</sup> |  |
|  | vB_OCPT_Car | 1.9 * 10<sup>9</sup> |  |
|  | vB_OCPT_Bop | 6.0 * 10<sup>7</sup> |  |
|  | vB_OCPT_Bill | 6.0 * 10<sup>7</sup> |  |
| ***Siphoviridae*** | vB_OCPT_SDS1 | 1.5 * 10<sup>8</sup> |  |
|  | vB_OCPT_SDS2 | 2.6 * 10<sup>7</sup> |  |
| ***Podoviridae*** | vB_OCPT_Ump | 1.6 * 10<sup>7</sup> |  |

D)
| Phage family | Phage name | Plaque Forming Unit (PFU/ml) | Colony Forming Unit of *E. faecalis* V587 (CFU/60µl) at OD<sub>600</sub> of 0.05 |
| --- | --- | --- | --- |
| ***Myoviridae*** | vB_OCPT_Carl | 1.3*10<sup>9</sup> | 7.2*10<sup>9</sup> |
|  | vB_OCPT_Ben | 1.3*10<sup>9</sup> |  |
|  | vB_OCPT_Car | 1.8*10<sup>8</sup> |  |
|  | vB_OCPT_Bop | 2.5*10<sup>7</sup> |  |
|  | vB_OCPT_Bill | 5*10<sup>9</sup> |  |
|  | vB_OCPT_Bob | 2.2*10<sup>9</sup> |  |
|  | vB_OCPT_V12 | 1.2*10<sup>9</sup> |  |

**Supplemental Table S2.**
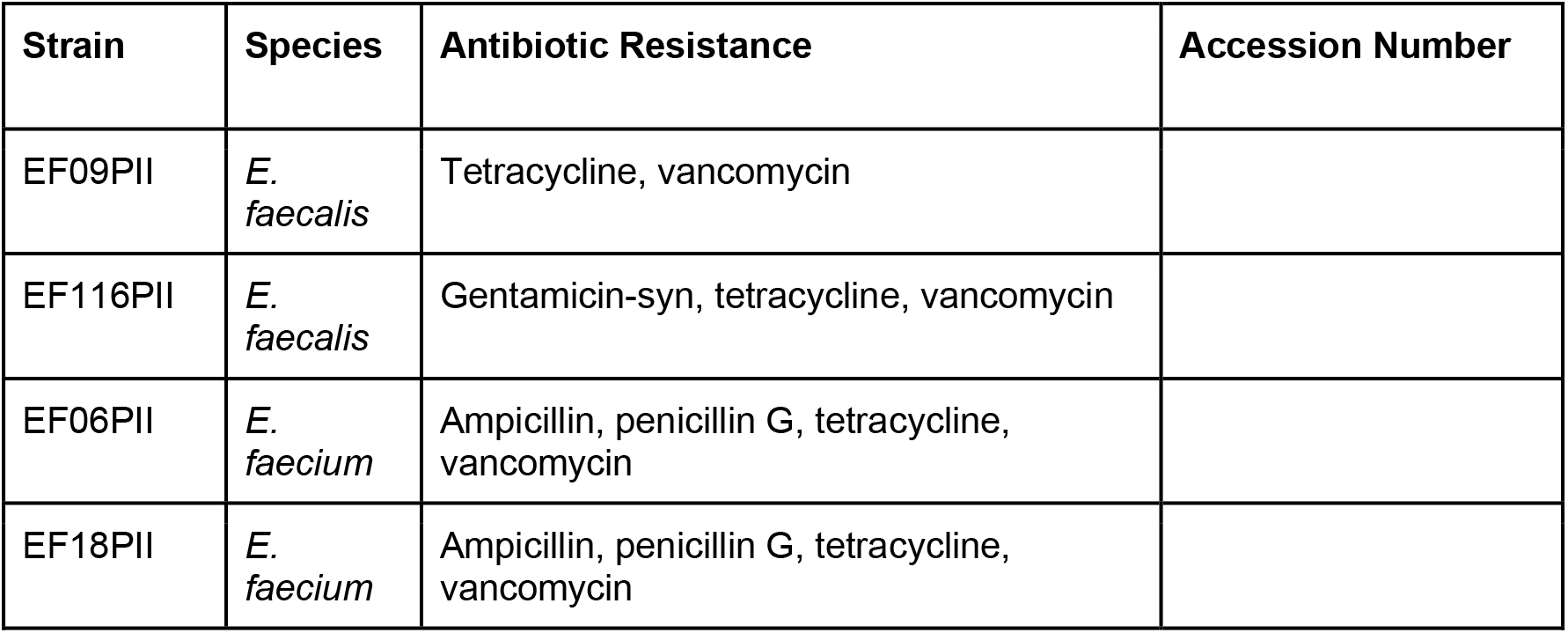
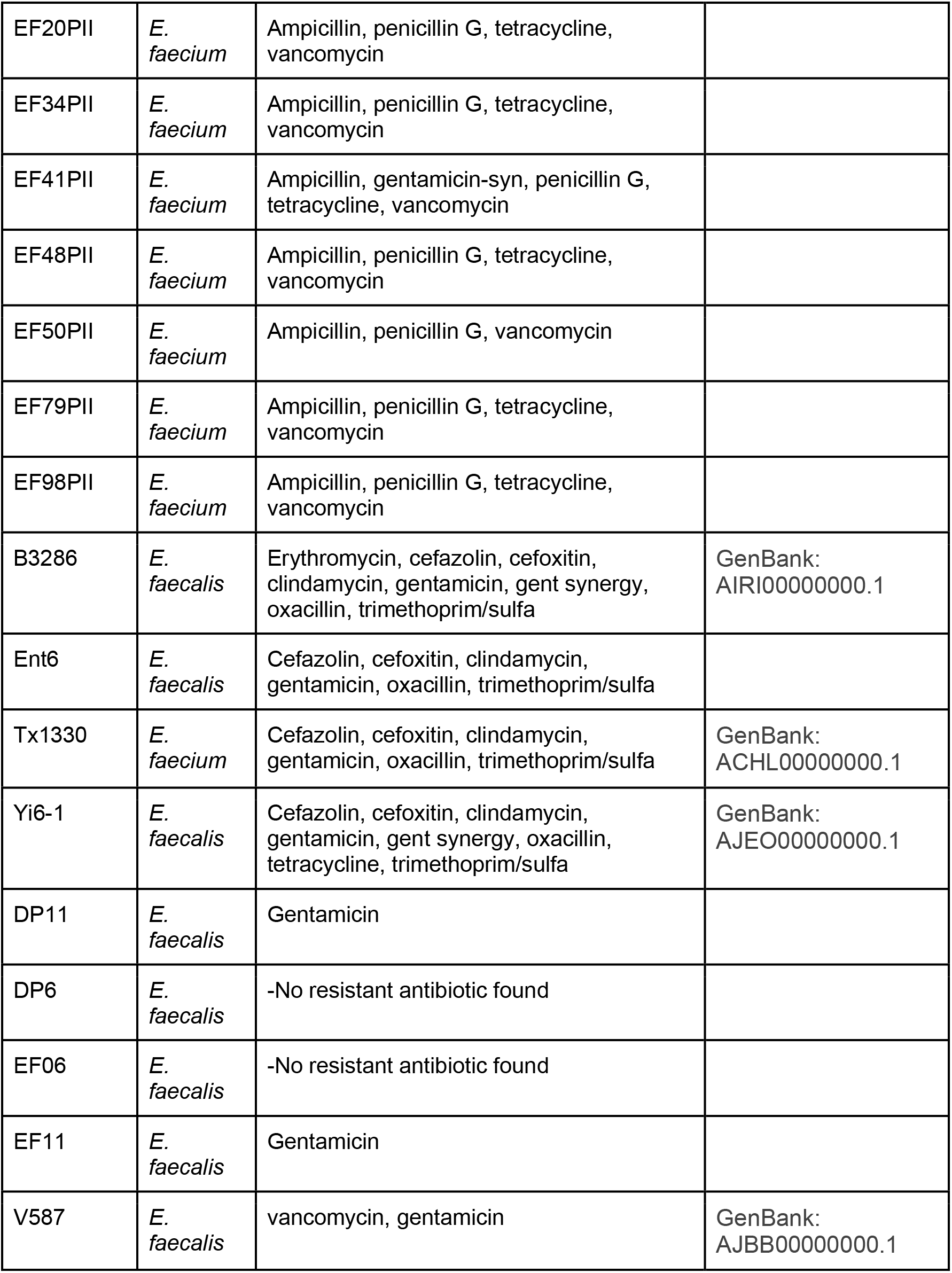
List of bacterial strains used in this study and the antibiotics to which they are resistant.

**Supplemental Table S3.** Phage information. The following phages were included in these experiments.

| Name | Family | Genus | Genome size |
| --- | --- | --- | --- |
| vB_OCPT_Bob | Myoviridae | Kochiodavirus | 150k |
| vB_OCPT_Car | Myoviridae | Kochiodavirus | 150k |
| vB_OCPT_Carl | Myoviridae | Kochiodavirus | 150k |
| EfV12-phi1 | Myoviridae | Schiekvirus | 150k |
| vB_OCPT_Ben | Myoviridae | Schiekvirus | 150k |
| vB_OCPT_Bop | Myoviridae | Shiekvirus | 150k |
| vB_OCPT_Bill | Myoviridae | Shiekvirus | 150k |
| vB_OCPT_CCS1 | Myoviridae | Shiekvirus | 150k |
| vB_OCPT_Tex | Myoviridae | Shiekvirus | 150k |
| vB_OCPT_SDS1 | Siphoviridae | Saphexavirus | 57k |
| vB_OCPT_SDS2 | Siphoviridae | Saphexavirus | 57k |
| vB_OCPT_CCS2 | Siphoviridae | Saphexavirus | 57k |
| vB_OCPT_CCS3 | Siphoviridae | Saphexavirus | 57k |
| vB_OCPT_Toy | Siphoviridae | Saphexavirus | 57k |
| vB_OCPT_CCS4 | Siphoviridae | Efquatrovirus | 40k |
| vB_OCPT_PG2 | Siphoviridae | Saphexavirus | 57k |
| vB_OCPT_PG9 | Siphoviridae | unclassified | 57k |
| vB_OCPT_PG11 | Siphoviridae | Saphexavirus | 57k |
| vB_OCPT_PG13 | Siphoviridae | Saphexavirus | 58k |
| vB_OCPT_Ump | Podoviridae | unclassified | 18k |

**Supplemental Table S4.**
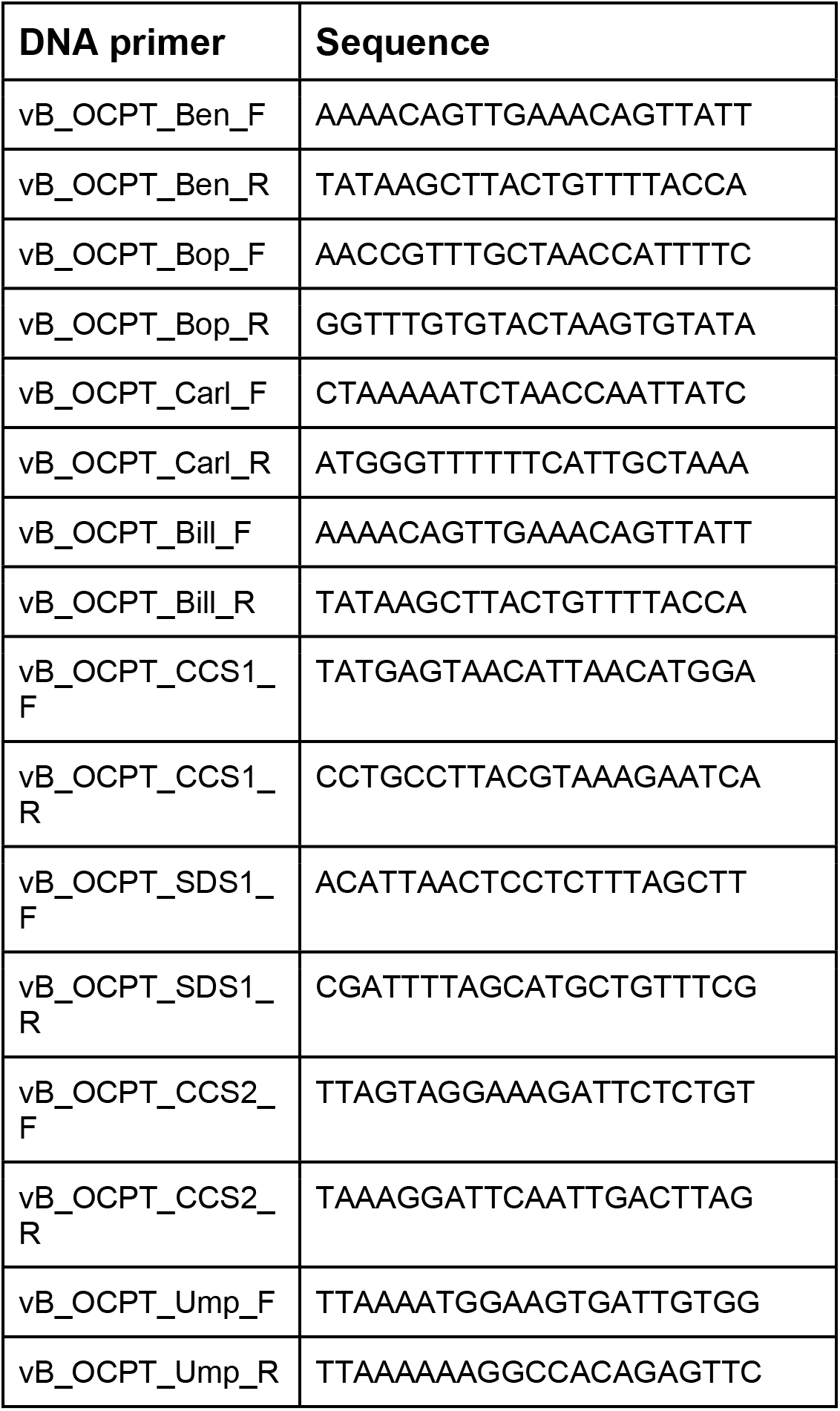
DNA primers (5’→3’) used to conduct qPCR experiments.

